# BeatRatePy - Automated High-Throughput Python-Based Beat Rate Quantification and Generation of Speed and Contraction Traces

**DOI:** 10.64898/2026.10.21.760713

**Authors:** Marie Held, William Partington, Thomas Waring, Aiswarya Dev, Lucy Isherwood, Brooke Boulton, Parveen Sharma, Rosalie Herbert, Tetsuhiro Kudoh, Alexander D. Corbett, Rebecca A.B. Burton

**Affiliations:** Centre for Cell Imaging, Technology, Infrastructure and Environment Directorate, Faculty of Health and Life Sciences, University of Liverpool, Liverpool, United Kingdom; Department of Pharmacology and Therapeutics, Institute of Systems, Molecular and Integrative Biology, University of Liverpool, Liverpool, United Kingdom; Cardiovascular and Metabolic Medicine, Institute of Life Course and Medical Sciences, University of Liverpool, Liverpool, United Kingdom; Biosciences, University of Exeter, Exeter, EX4 4QD, United Kingdom; Department of Physics and Astronomy, University of Exeter, EX4 4QL, United Kingdom; Living Systems Institute, University of Exeter, EX4 4QD, United Kingdom; Liverpool Centre for Cardiovascular Science at University of Liverpool, Liverpool John Moores University and Liverpool and Heart and Chest Hospital, Liverpool, United Kingdom; Department of Pharmacology, University of Oxford, Oxford, United Kingdom

**Keywords:** Cardiomyocyte contraction analysis, Label-free imaging, Beat rate quantification, Pixel variance analysis, High-throughput cardiac imaging

## Abstract

Analysing cardiac motion from label free imaging can be difficult, especially when recordings vary in quality or show complex movement. We developed a Python-based pipeline that automatically measures beat rate, contraction, speed from transmitted light timelapse data. The workflow runs in a Jupyter notebook and uses straightforward steps that can be followed easily, making the analysis accessible including to users with limited coding experience. Using human iPSC-derived cardiomyocytes imaged on a Zeiss Cell Discoverer 7, the pipeline produced beat rates, contraction and contraction speed measurements that matched expected behaviour. To test wider use, we applied the same workflow to killifish larvae heart recordings and public datasets, changing only basic input parameters. The pipeline performed consistently across these different systems and did not require changes to the underlying code. By supporting clear parameter control and use across laboratories, microscopy platforms and repositories, this approach strengthens reproducible analysis of cardiac motion from label free imaging.

**NEW & NOTEWORTHY:** We have established a protocol for label-free transmitted light imaging on a Zeiss Cell Discoverer 7 platform which enables contraction visualisation and quantification of human iPSC-derived cardiomyocytes (hiPSC-CMs). This provides the input for a new, user-friendly Python script based on the Pixel Variance algorithm, that has been developed to enable automatic batch processing of recordings to analyse beat rate and contraction.

## INTRODUCTION

Cardiac tissue models are essential for drug screening, understanding developmental biology and pathophysiological analysis^1^. The *in vivo* cardiac models are some of the most basic, using small and large animals such as rodents and pigs to study underlying mechanisms of cardiovascular disease (CVD) as well as determining safety and efficacy thresholds for pharmaceuticals that can be converted into equivalent human doses^2,3^. Many of these small animal *in vivo* models struggle to accurately translate into human activity due to a lack of complexity, whilst larger and more accurate models like pigs provide a greater ethical challenge and increased costs^4^. *In vitro* model alternatives are much more cost effective and easier to work with, such as zebrafish primary cardiomyocytes (CMs), neonatal rat CMs and mouse CMs^4-6^. Whilst these cell lines maintain similar phenotypes to those *in vivo*, they have a limited lifespan that restricts the time frame over which experiments can be conducted^4^. Immortalised human cardiac cell lines like the AC16 line provides an ethically simple and easily cultured alternative by retaining cardiac molecular features over a sustained period of time, however a lack of contractility makes it inappropriate to study wave dynamics or arrhythmias; this means animal models still have an important role^7,8^.

Stem cell-based models have been predicted as the future of medical research, offering a wide range of cell types, that can be grown in a laboratory, which possess relatively high genotypic accuracy^9-12^. The use of human induced pluripotent stem cell-derived cardiomyocytes (hiPSC-CMs) in mechanistic studies, drug development and cardiotoxicity models is becoming more commonplace, with the longer lifespan of the model suiting it to test multiple-exposure effects^13-16^. These models can be assessed and quantified by a variety of methods, primarily involving electrophysiology analysis, high-throughput screens and cell imaging^17-21^. Another commonly used model for studying cardiac development, ageing and regeneration^22^ is killifish larvae. Killifish are the shortest-lived vertebrate, allowing it to be used as a model for ageing.

Imaging is a frequently employed technique to study the functionality of CMs non-invasively. The use of calcium and voltage dyes are commonplace in CM imaging to produce robust functional readout data^23,24^. By using calcium-sensitive fluorescent dyes, the location and concentration of calcium ions can be detected during imaging and subsequently, action potential duration can be monitored and the presence of arrhythmias detected^21,25^. However, both voltage and calcium dyes possess inherent toxic effects that limit exposure and make them inappropriate for long term studies^23,26^. Without careful selection of the correct dyes and optimisation of experimental protocols, dyes have the potential to alter the physiology and pharmacological response of hiPSC-CMs^23^.

It is therefore preferable to exclude dyes from imaging whilst retaining the robust data and image analysis opportunity it provides. One alternative is to use label-free imaging and rely on post-image processing to identify wave dynamics^27^. Liu et al.,^27^ found that the non-invasive dye-free alternatives matched and in parts surpassed the sensitivity of dye-based imaging. Other dye-free methods involve using optogenetic stimulation to initiate contraction of cardiac waves and then image them using off-axis oblique illumination, as well as other phase-contrast imaging methods^25,26^. To achieve accurate long-term imaging of hiPSC-CMs, microscopes must accurately control temperature and gas similar to that of an incubator, as well as provide high speed recording capabilities to allow post-image processing to differentiate between different waves.

One such microscope is the Zeiss Cell Discoverer 7 (CD7), a flexible imaging platform which can be used to record a wide range of subjects from long-term live epifluorescence imaging of cricket embryos to hiPSC-derived neurons^28,29^. The inbuilt incubation capabilities make it suitable for imaging samples over prolonged periods of time, and is easy to use partly due to its autofocus capabilities and detection of the thickness and optical properties of the sample carrier^30^. The CD7 can also be set up to automatically acquire images regularly in specific regions of interest at designated time intervals, which, along with its capability of recording high-speed time series, makes it suitable for highly dynamic processes. Multiple transmitted light options in the CD7 enable label-free imaging of live, transparent cells, enhancing visibility of structures like cell boundaries, nuclei, vacuoles, organelles and therefore tracking cell movement. The instrument provides a fully enclosed and mechanically stable optical path that minimises drift and vibration, ensuring that time-series recordings remain spatially consistent even when the sample itself is moving. It offers high-quality transmitted-light contrast modes which generate sharp, high-contrast visualisation of cell boundaries, and contraction dynamics without the need for fluorescent labels. This reduces phototoxicity and avoids the physiological perturbations associated with calcium dyes or other reporters. Illumination is LED-based, uniform and photometrically stable, supporting quantitative analysis of motion, wave fronts or deformation fields in transmitted light. In addition, the CD7 is built for automation: its plate-handling capabilities, reproducible stage movement and software-integrated workflows allow systematic sampling of multiple wells and fields, which is particularly useful for pharmacological testing, high-content screening or longitudinal functional assays. Together, these features create an imaging environment that combines low phototoxic load, high contrast, temporal stability and robust autofocus, enabling accurate, long-duration recordings of cardiomyocyte behaviour without perturbing the cells.

In this study, we utilised hiPSC-CMs and performed cardiac imaging on the CD7 microscope. The high-quality image files taken with the CD7 were collected in the ZEN Blue control software and saved as CZI files, typically 4 to 6 GB in size for a 20-second timelapse. A single experiment using 20 wells and 2 or 3 time points generates 40-60 time lapse files for analysis. To demonstrate the versatility of BeatRatePy, label-free killifish data was also taken using handheld digital microscopes at frame rates of 5-10 Hz.

Python is a coding language through which custom scripts can be written and run which also enables batch processing of multiple, large time lapse files. We aimed to create and fine-tune a rapid analysis pipeline to quantify beat rate, speed and contraction dynamics from long-term transmitted-light imaging either as single time lapse files or in batch. In addition, we analysed killifish larvae heartbeat rate as well as some publicly available datasets and demonstrate wider applicability of our script. The python script is implemented in a Jupyter notebook for user-friendliness and ease of use. BeatRatePy was designed to address several limitations of existing tools for label-free cardiac motion analysis.

## RESULTS

BeatRatePy calculates the beat rate from imaged cardiac monolayers, spheroids and beyond, imaged in transmitted light mode on a high throughput microscope (Table 1). It is set up to import the proprietary microscope image file formats.

**Table 1:**
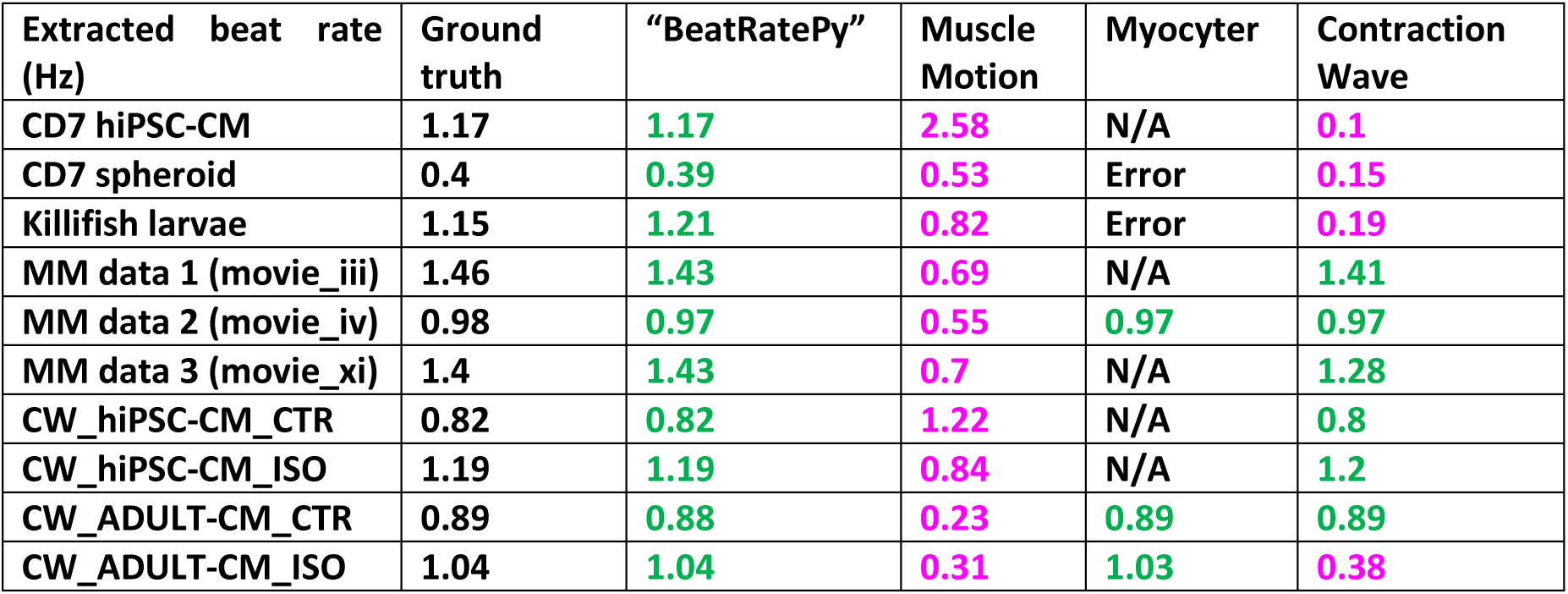
Summary of results extracted for each of the analysis software packages considered. **Note:** Myocyter: N/A = could not be analysed as not a single cell; Error = processing resulted in an error message, used the recommended Fiji version (2.0.0-rc-69/1.52n).

| Extracted beat rate (Hz) | Ground truth | "BeatRatePy" | Muscle Motion | Myocyter | Contraction Wave |
| --- | --- | --- | --- | --- | --- |
| CD7 hiPSC-CM | 1.17 | 1.17 | 2.58 | N/A | 0.1 |
| CD7 spheroid | 0.4 | 0.39 | 0.53 | Error | 0.15 |
| Killifish larvae | 1.15 | 1.21 | 0.82 | Error | 0.19 |
| MM data 1 (movie_iii) | 1.46 | 1.43 | 0.69 | N/A | 1.41 |
| MM data 2 (movie_iv) | 0.98 | 0.97 | 0.55 | 0.97 | 0.97 |
| MM data 3 (movie_xi) | 1.4 | 1.43 | 0.7 | N/A | 1.28 |
| CW_hiPSC-CM_CTR | 0.82 | 0.82 | 1.22 | N/A | 0.8 |
| CW_hiPSC-CM_ISO | 1.19 | 1.19 | 0.84 | N/A | 1.2 |
| CW_ADULT-CM_CTR | 0.89 | 0.88 | 0.23 | 0.89 | 0.89 |
| CW_ADULT-CM_ISO | 1.04 | 1.04 | 0.31 | 1.03 | 0.38 |

### hiPSC-CM Cultures

To validate BeatRatePy, we first applied it to our own dataset of hiPSC-CMs, cultured and imaged using the established label-free transmitted light protocol on the Zeiss Cell Discoverer 7 platform (Table 2, Figure 1). This dataset served as the primary benchmark during translation of the original Matlab code (used in^31^) into Python. The pipeline successfully extracted contraction waveforms, beat-to-beat contraction traces, and enabling automated quantification of contraction amplitude and beat frequency across multiple recordings.

**Figure 1:**
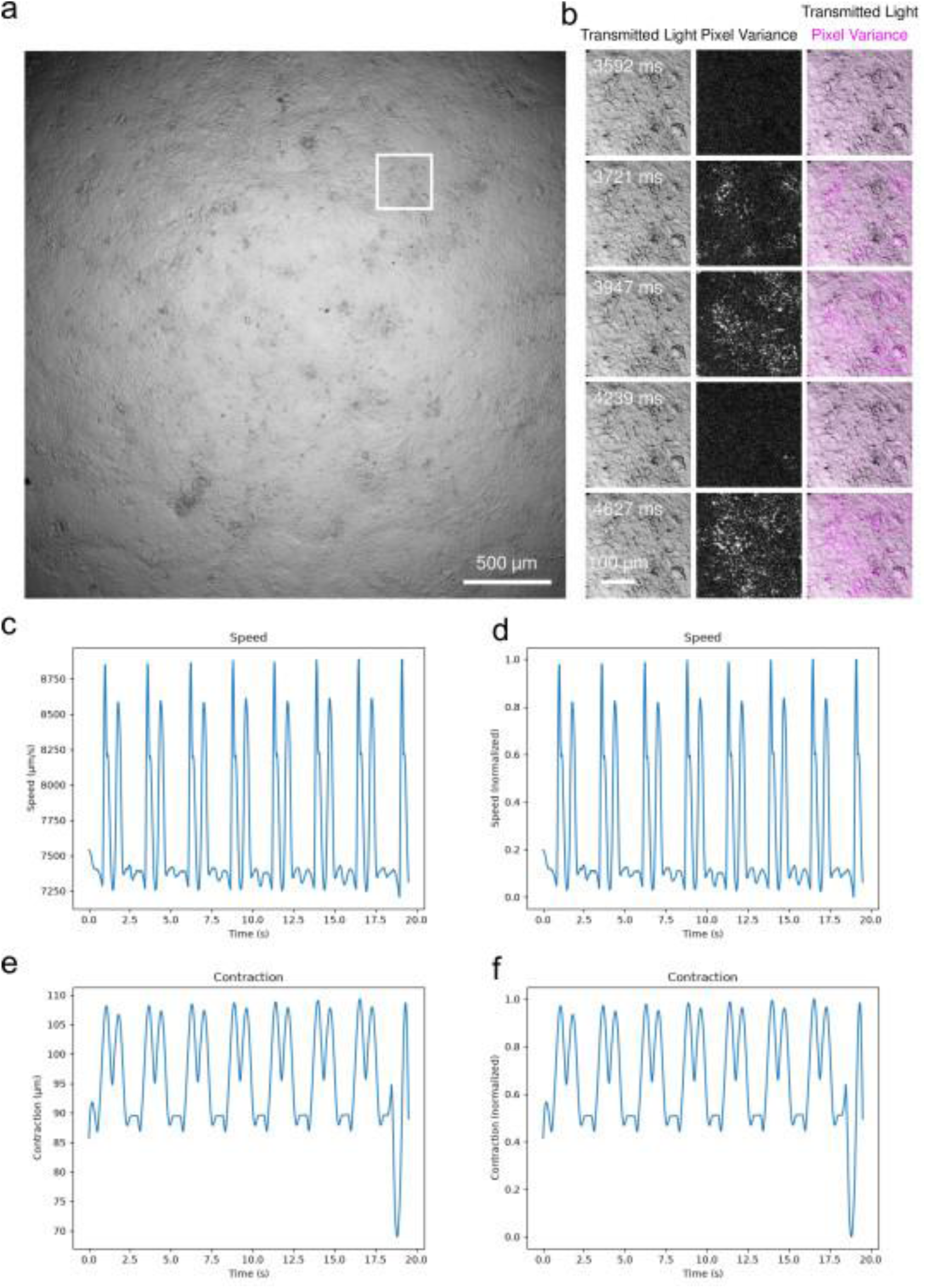
Analysis of a spontaneously beating hiPSC-cardiomyocyte monolayer imaged by transmitted-light microscopy on the Zeiss Cell Discoverer 7 (CD7). a) whole field of view captured. b) selected time points from the time-lapse recording; the original field of view was cropped to highlight a region of contractile activity (solid line inset frame shown in a). For each time point, the original input image, the corresponding pixel variance image, and a merged image showing the pixel variance signal overlaid on the original image are displayed. c-f) speed and contraction traces generated by BeatRatePy, shown as scaled and normalised outputs. The pixel variance approach enables visualisation and quantification of cyclic cardiomyocyte motion and deformation over time^39^. See Supplementary Video 1 and 2 for the complete time series.

**Table 2:** Microscope components and acquisition parameters used for imaging on Zeiss Cell Discoverer 7. Comparison of settings used for transmitted-light time-series imaging of hiPSC-cardiomyocyte monolayers, and transmitted-light time-series imaging of cardiac spheroids. Parameters reported include illumination settings, optical configuration, objective lens characteristics, detector specifications, exposure times, image bit depth, and spatial calibration. These acquisition settings were used to generate the image datasets subsequently analysed using BeatRatePy pipeline.

| Microscope component | Parameters for cell culture time series | Parameters for cardiac spheroid time series |
| --- | --- | --- |
| Light source | White light LED lamp<br>1% intensity | White light LED lamp<br>12% intensity |
| Excitation/Emission optics | Contrast method: Transmitted light - Brightfield<br>Filters: 412-438, 501-527, 583-601, 662-756<br>Beam Splitter: 394, 490, 573, 691<br>Filter Emission Wavelength: 412-438, 501-527, 583-601, 662-756 | Contrast method: Transmitted light - Brightfield<br>Filters: 412-438, 501-527, 583-601, 662-756<br>Beam Splitter: 394, 490, 573, 691<br>Filter Emission Wavelength: 412-438, 501-527, 583-601, 662-756 |
| Objective lens | Plan-Achromat 5×/ 0.35 NA<br>Depth of focus: 8.98 μm<br>With a 0.5× tube lens<br>Effective NA: 0.125 | Plan-Achromat 5×/ 0.35 NA<br>Depth of focus: 8.98 μm<br>With a 0.5× tube lens<br>Effective NA: 0.125 |
| Detector | Zeiss AxioCam 712 mono<br>1× camera adapter<br>Exposure time: 619 μs<br>Binning: 1, 1<br>Bit depth: 8 bit<br>Sampling: 1.38 μm / pixel | Zeiss AxioCam 712 mono<br>1× camera adapter<br>Exposure time: 200 μs<br>Binning: 1, 1<br>Bit depth: 14 bit<br>Sampling: 1.38 μm/pixel |
| Time-lapse | Interval: 619 μs<br>Number of frames: 612<br>Total Duration: 19.77 sec | Interval: 200 μs<br>Number of Frames: 2530<br>Total duration: 19.69 sec |

Each time-lapse contained a single, uncropped, beating monolayer. The input parameters frame rate and pixel calibration were taken directly from CD7 metadata. For ground truth data generation, the beat rate was estimated manually by counting contractions over the recording. The frame offset was calculated from the frame rate, measured beat rate, and an offset fraction of 0.2 (see ‘Time Series Analysis’ below). The quantitative outputs aligned well with expected physiological behaviour of hiPSC-CMs. Contraction frequencies fell within the typical spontaneous beating range for immature CMs, and waveform morphology showed the characteristic rapid upstroke and slower relaxation phase. These findings confirm that the pipeline captures biologically meaningful contractile dynamics rather than artefactual motion.

Together, this validation demonstrates that the Python framework is not only technically robust but also faithfully reflects known features of human CM behaviour, supporting its use for high-throughput, label-free analysis of cardiac dynamics.

### Spheroid

Cardiac spheroids were also imaged using the established label-free transmitted light protocol on the Zeiss Cell Discoverer 7 platform (Figure 2). The input parameters frame rate and pixel calibration were taken directly from CD7 metadata, the ground truth beat rate was estimated manually by counting contractions over the recording, and the frame offset was calculated from the frame rate, measured beat rate, and an offset fraction of 0.2. Where multiple spheroids were present in the same field of view, individual spheroids were cropped prior to analysis (see Materials and Methods, Data Pre-processing) (Figure 2).

**Figure 2:**
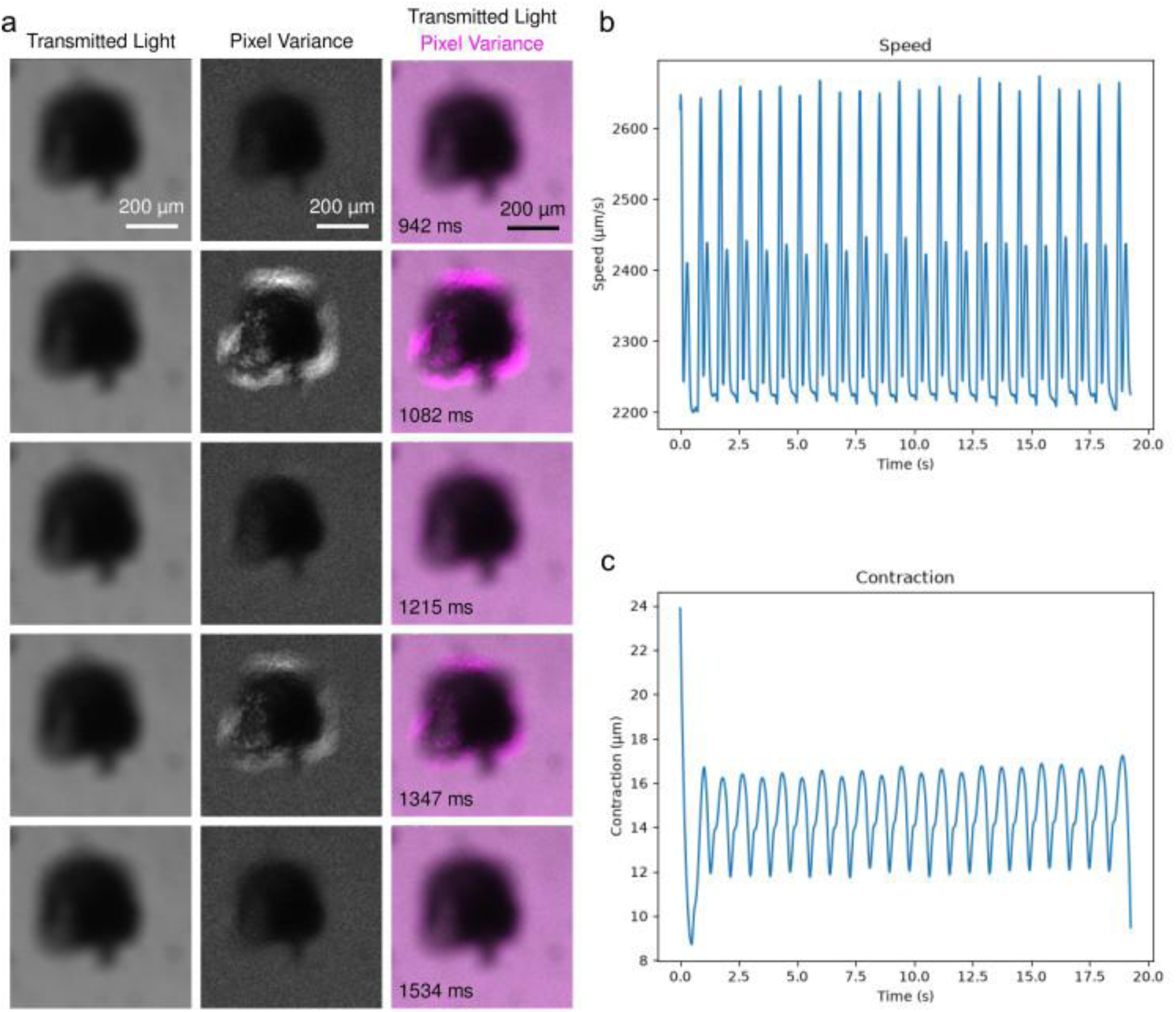
Analysis of a spontaneously beating hiPSC-cardiac spheroid imaged by transmitted-light microscopy on the Zeiss Cell Discoverer 7 (CD7). Individual spheroids were cropped from the original field of view prior to analysis when multiple spheroids were present within the recording. a) selected time points from the transmitted-light time-lapse sequence showing the cardiac spheroid during successive contraction and relaxation phases. For each time point, the original input image, the corresponding pixel variance image, and a merged image displaying the pixel variance signal overlaid on the original image are shown. b & c) speed and contraction traces generated by BeatRatePy, presented as scaled outputs. The pixel variance approach enables visualisation and quantification of cyclic motion and deformation within the spheroid, allowing automated assessment of contractile activity and beat frequency. See Supplementary Video 3 for the complete time series.

### Killifish

To evaluate the flexibility of BeatRatePy, we subsequently applied it to cardiac imaging data from killifish, a commonly used small-vertebrate model^32^. While the imaging approach and acquisition parameters are detailed in the Materials and Methods, this dataset differs markedly from hiPSC-CM monolayers in both spatial scale and motion dynamics. Whole-heart contractions produce larger displacements, multi-axis deformation, and greater regional heterogeneity.

Killifish cardiac videos were converted from AVI to TIFF stacks using FFMPEG and Fiji, with frames transformed from RGB to 8-bit grayscale. Each recording contained multiple embryos and was cropped to isolate the developing, beating heart. Input parameters were derived from the video metadata: frame rate was extracted via FFMPEG, ground truth beat rate was estimated by analysing intensity fluctuations over time in Fiji, and pixel calibration was not available for this dataset. The frame offset was automatically calculated from the frame rate, the measured beat rate, and an offset fraction of 0.2.

Employing the same analysis pipeline with only minor, dataset-specific parameter adjustments, the framework reliably extracted sped and contraction profiles, and beat-to-beat signals without any need for structural changes to the codebase (Figure 3). The resulting quantitative measurements were consistent with established killifish cardiac physiology, including increased contraction magnitude, more rapid ventricular propagation, and stronger regional variation relative to cultured cardiomyocytes^32^. The measured beat rate of 1.21 Hz is in good agreement with the ground truth value of 1.15 Hz (Table 1).

**Figure 3:**
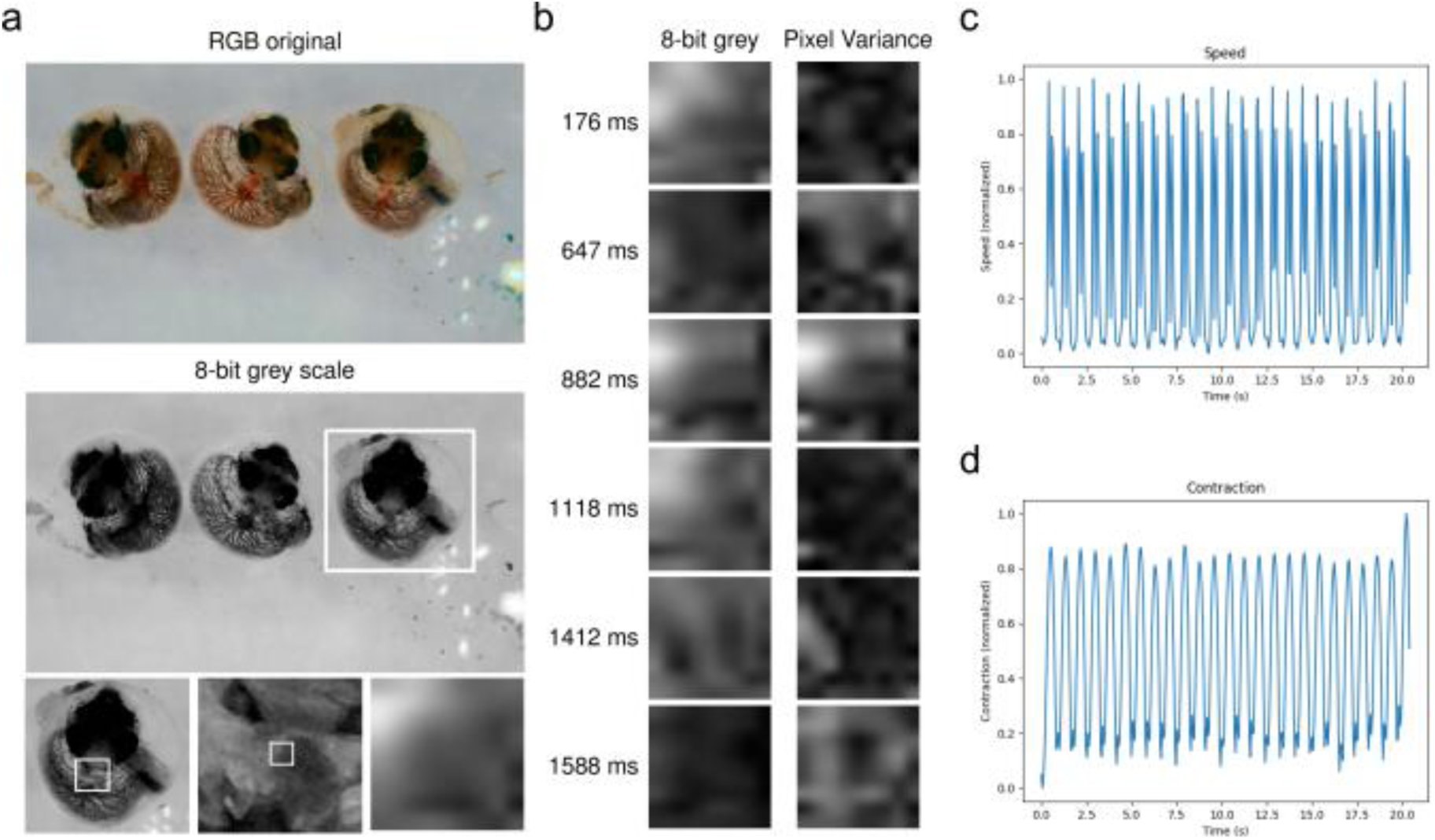
Analysis of a killifish heart recording using BeatRatePy. Data obtained from preprint^32^ a) original colour video frame (top), corresponding 8-bit grayscale image (bottom), and progressive inset regions illustrating the sequential cropping and zooming steps used to isolate the beating heart. B) selected time points from the processed video; the left column shows the final cropped grayscale input images and the right column shows the corresponding pixel variance images generated by BeatRatePy. c & d) speed trace calculated from the first 375 frames of the recording, demonstrating periodic cardiac contractions. Scale information was not available for the original dataset.

### MUSCLEMOTION - MM Data set 1 (movie iii)

MM Data set 1 (movie iii) comprises a monolayer of a beating, contracting hiPSC-CM culture recorded in brightfield transmitted light mode. The beat rates measured by BeatRatePy (1.43 Hz) and by Contraction Wave (1.41 Hz) were in good agreement with the manually determined ground truth (1.46 Hz), while the original publication reported that the culture was paced at 1.5 Hz (Table 1). By contrast, the beat rate calculated from MUSCLEMOTION peak positions was only around half of the ground truth value; beat rate is not a parameter that can be extracted directly from the MUSCLEMOTION macro (Figure 4).

**Figure 4:**
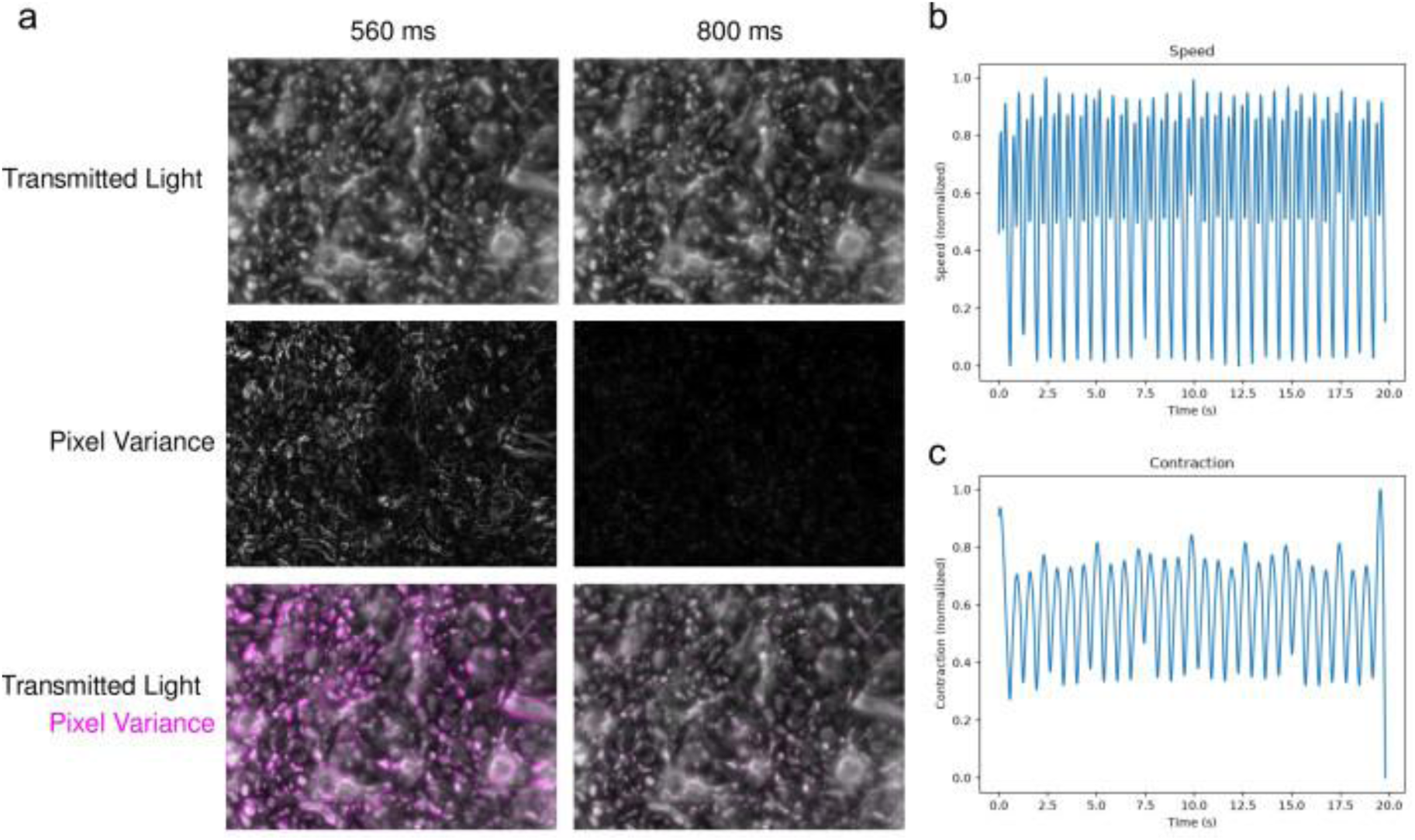
Analysis of MM Data set 1 (movie iii) using BeatRatePy. Representative analysis of a transmitted light phase contrast recording of a contracting human stem cell-derived cardiomyocyte monolayer. a) Multi-panel image showing selected frames, at two different time points, from the original time-lapse recording to illustrate periodic contraction of the culture (top), as well as the corresponding pixel variance images generated by BeatRatePy highlighting regions of motion between successive frames (middle) and the merge of both images (bottom). b) normalised speed and contraction traces calculated from the pixel variance analysis. Regular peaks correspond to individual contraction cycles and were used for automated beat-rate determination. BeatRatePy measured a beat rate of 1.43 Hz, closely matching both the manually determined ground truth (1.46 Hz) and the reported pacing frequency (1.5 Hz). Scale information was not available from the original published dataset. Original data from^33^.

### MUSCLEMOTION - MM Data set 2 (movie iv)

MM Data set 2 (movie iv) comprises a beating, contracting human stem cell-derived cardiac spheroid recorded in phase contrast mode. The beat rates measured by BeatRatePy (0.97 Hz) and by Contraction Wave (0.97 Hz) were in good agreement with the ground truth (0.98 Hz), whereas the beat rate calculated from MUSCLEMOTION peak positions was substantially lower, at 0.55 Hz (Table 1, Figure 5).

**Figure 5:**
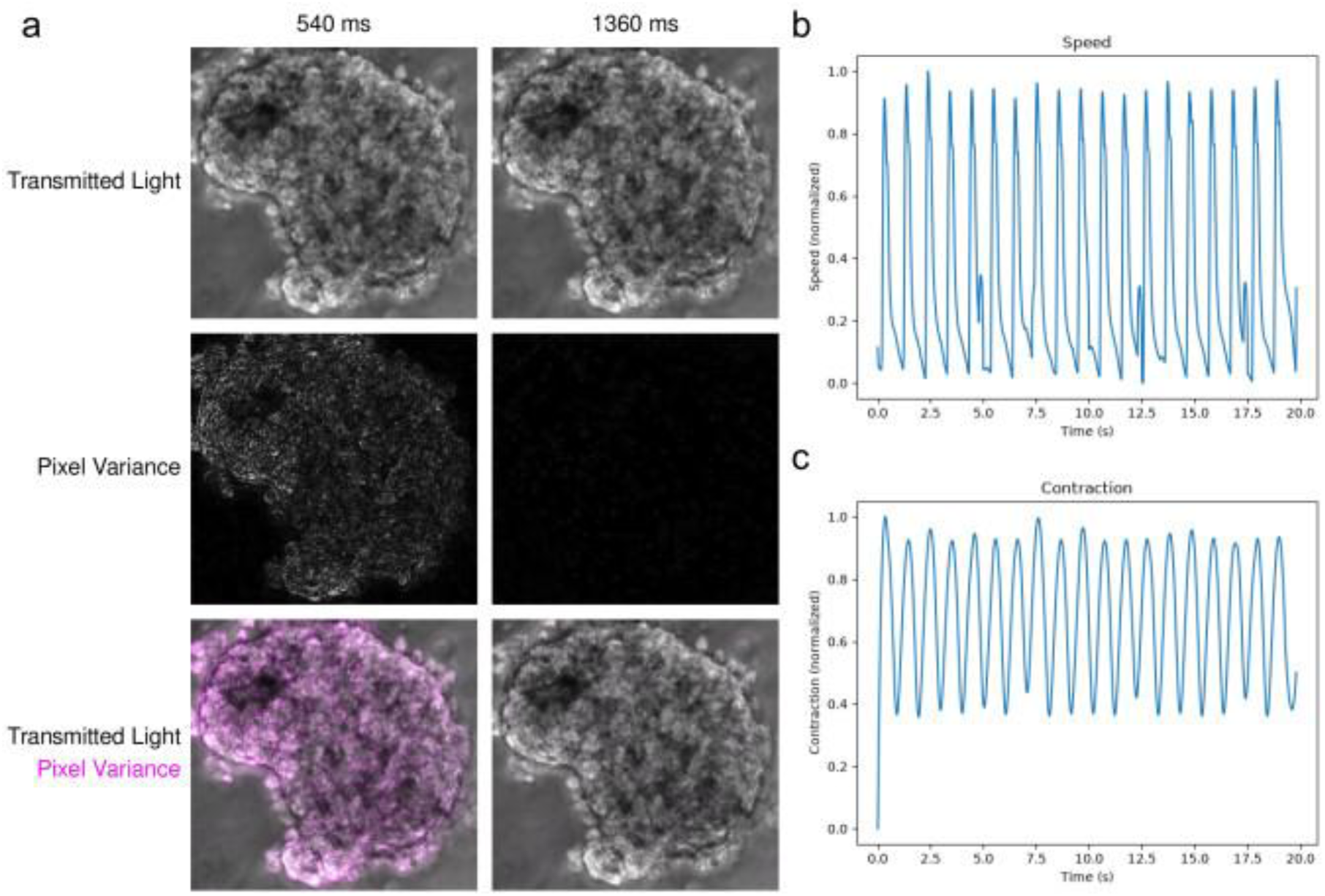
Analysis of MM Data set 2 (movie iv) using BeatRatePy. a) selected time points from a transmitted light phase-contrast recording of a beating human stem cell-derived cardiac spheroid (top), as well as the corresponding pixel variance images generated by BeatRatePy highlighting regions of motion during contraction and relaxation (middle) and the merge of both images (bottom). b) calculated, normalised speed and contraction traces. The periodic peaks detected in the traces were used for automated beat-rate determination. BeatRatePy measured a beat rate of 0.97 Hz, in close agreement with the manually determined ground truth value of 0.98 Hz. Scale information was not available from the original published dataset. Original data from^33^.

### MUSCLEMOTION - MM data 3 (movie_xi)

MM Data set 3 (movie xi) is a human echocardiogram recording of a healthy individual. The beat rate calculated by BeatRatePy (1.43 Hz) was in good agreement with the ground truth (1.4 Hz), whereas Contraction Wave and MUSCLEMOTION underestimated the beat rate by 9% and 50%, respectively (Table 1, Figure 6).

**Figure 6:**
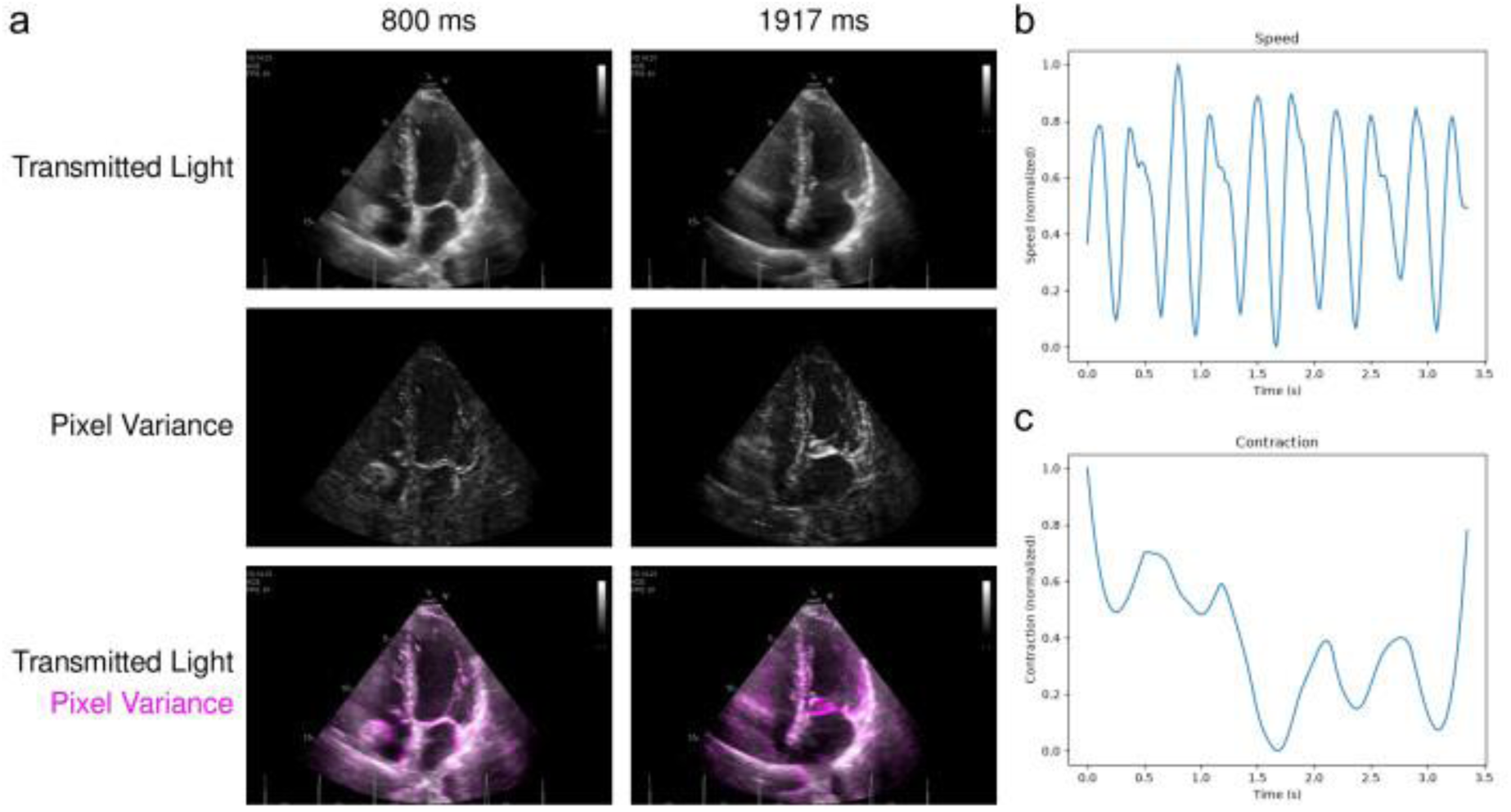
Analysis of MM Data set 3 (movie xi) using BeatRatePy. a) Multi-panel image showing selected time points from a human echocardiogram recording of a healthy individual obtained from the public MUSCLEMOTION dataset^33^ (top), as well as corresponding pixel variance images generated by BeatRatePy highlighting regions undergoing cyclic motion during the cardiac cycle (middle) and the merged images (bottom). b): calculated, normalised speed and contraction traces. Regular oscillations in the traces enabled automated beat-rate determination. BeatRatePy measured a beat rate of 1.43 Hz, in close agreement with the manually determined ground truth value of 1.40 Hz, demonstrating applicability of the pipeline to clinical cardiac imaging data beyond cell-based preparations. Scale information was not available from the original published dataset. Original data from^33^.

### Contraction Wave (CW) Data set 1 & 2 - CW_hiPSC-CM_CTR and CW_hiPSC-CM_ISO

For the CW_hiPSC-CM_CTR and CW_hiPSC-CM_ISO datasets, beat rates calculated by BeatRatePy were in perfect agreement with the ground truth for both the control (0.82 Hz) and isoproterenol (ISO)-treated (1.19 Hz) cultures. ISO treatment increased the contractility of the culture, clearly visible in the speed and contraction traces, and the drug-induced differences were clearly identified by the pipeline. MUSCLEMOTION overestimated the beat rate of the control culture (1.22 Hz) and underestimated that of the ISO-treated culture (0.84 Hz), while Contraction Wave calculations were in good agreement with the ground truth; 0.82 Hz and 1.19 Hz for the control and treated culture respectively (Table 1, Figure 7).

**Figure 7:**
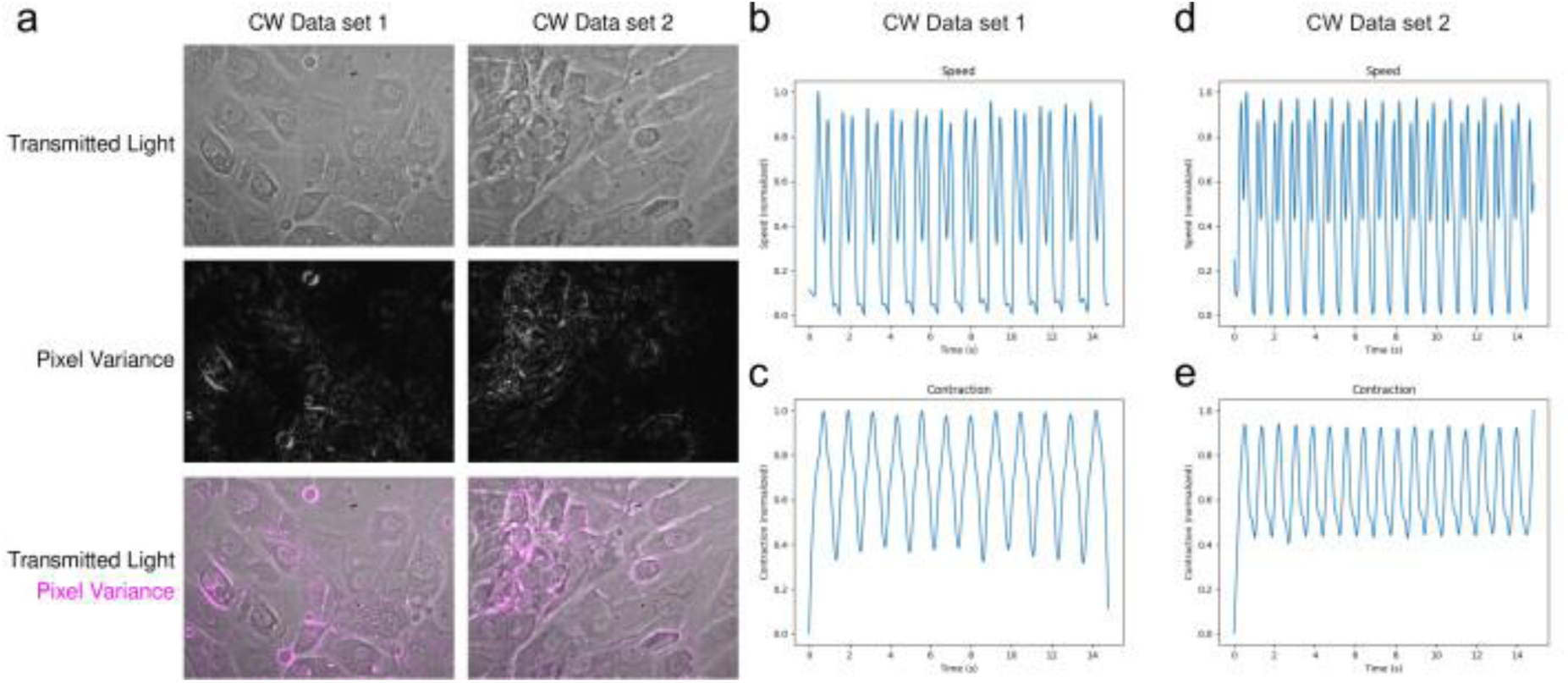
Analysis of the ContractionWave datasets CW_hiPSC-CM_CTR and CW_hiPSC-CM_ISO using BeatRatePy. a) selected time points from transmitted-light recordings of human induced pluripotent stem cell-derived cardiomyocyte (hiPSC-CM) cultures under control conditions (CTR) and following isoproterenol (ISO) treatment (top), as well as corresponding pixel variance images generated by BeatRatePy highlighting regions of contractile motion (middle) and the merge of both images (bottom). :b-e) calculated, normalised speed and contraction traces. Isoproterenol treatment increased spontaneous beating frequency and altered contraction dynamics, evident from the increased frequency of trace oscillations. Automated beat-rate analysis produced values of 0.82 Hz and 1.19 Hz for the control and isoproterenol-treated cultures, respectively, in agreement with the manually determined ground truth values. Scale information was not available from the original dataset. Original data from^33^.

### Contraction Wave (CW) Data set 3 & 4 – CW_ADULT-CM_CTR and CW_ADULT-CM_ISO

For the CW_ADULT-CM_CTR and CW_ADULT-CM_ISO datasets (Table 3), beat rates calculated by BeatRatePy (control: 0.88 Hz; isoproterenol-treated: 1.04 Hz) were in very good agreement with the ground truth (control: 0.89 Hz; isoproterenol-treated: 1.04 Hz) (Table 1, Figure 8). Isoproterenol treatment increased the contractility of the culture, clearly visible in the speed and contraction traces, and the drug-induced differences were clearly identified. MUSCLEMOTION severely underestimated the beat rate of both the control (0.23 Hz) and isoproterenol-treated (0.31 Hz) cultures. Contraction Wave calculations were in good agreement with the ground truth for the control condition (0.89 Hz) but not for the treated condition (0.38 Hz).

**Figure 8:**
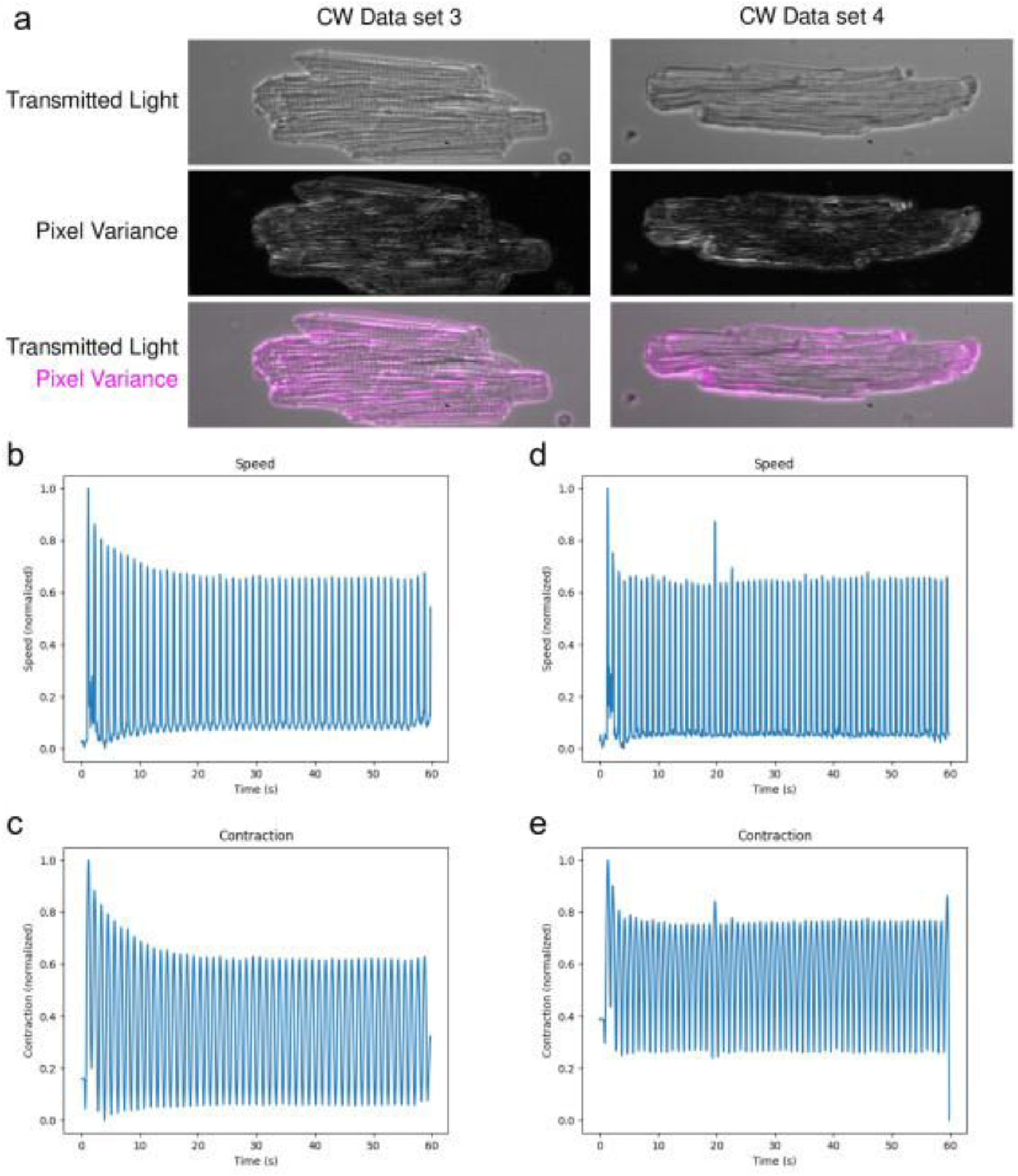
Analysis of the ContractionWave (CW) datasets in adult cardiomyocytes (CM), CW_ADULT-CM_CTR and CW_ADULT-CM_ISO using BeatRatePy. a) selected time points from recordings of acutely isolated adult ventricular cardiomyocytes under control conditions (CTR) and following isoproterenol (ISO) treatment (top), as well as corresponding pixel variance images generated by BeatRatePy highlighting regions of contractile motion throughout the contraction cycle (middle) and the merge of both images (bottom). b-e) calculated, normalised speed and contraction traces. Isoproterenol (ISO) treatment increased beating frequency and altered contractile behaviour, reflected by changes in the temporal profiles of the speed and contraction traces. Automated beat-rate analysis yielded values of 0.88 Hz for control cells and 1.04 Hz for ISO-treated cells, in close agreement with the manually determined ground truth values of 0.89 Hz and 1.04 Hz, respectively. Scale information was not available from the original published dataset. Original data from^33^.

## DISCUSSION

This work introduces a versatile and user-friendly pipeline for analysing label-free cardiac timelapse recordings, delivered through a Jupyter notebook to enhance clarity and ease of use. The same analytical workflow operated effectively across hiPSC-CMs, killifish heart recordings, and publicly available datasets^32^, demonstrating strong performance despite substantial differences in scale, motion patterns, and signal quality. This breadth of applicability contrasts with many existing cardiac-motion analysis tools, which are often restricted to specific experimental setups or rely heavily on manual processing.

Building on the earlier MATLAB implementation, the Jupyter-based version offers a more accessible and transparent environment in which all parameters can be viewed and modified directly. This design aligns with FAIR principles and supports straightforward reuse across research groups and transmitted light imaging modalities. Because the pipeline is built entirely within the open Python ecosystem, it avoids the limitations of proprietary or paywalled software. It also complements tools such as MUSCLEMOTION^33-35^, Myocyter and ContractionWave by providing a label-free approach optimised for transmitted-light imaging and capable of extracting both contraction and wave-propagation metrics from raw proprietary imaging data files.

While the datasets analysed here primarily serve to demonstrate feasibility rather than uncover new biology, the pipeline provides robust quantitative measures of contractile behaviour that can underpin future mechanistic studies. Metrics such as contraction amplitude, beat rate, and propagation dynamics offer a foundation for comparing developmental stages, disease models, or pharmacological responses.

In summary, this work delivers a reliable, broadly applicable, accessible and extensible platform for analysing cardiac dynamics from diverse imaging modalities, reducing technical barriers and strengthening reproducibility in functional cardiac imaging research.

## LIMITATIONS

The performance of the pipeline remains dependent on the quality and stability of the input recordings. Suboptimal imaging conditions such as low contrast, illumination fluctuations, high noise or sample drift, may reduce the accuracy of motion and contraction estimates. Only some of these can be addressed via image pre-processing. Extreme or irregular movements can also challenge the pixel-variance approach, occasionally leading to incomplete or noisy traces. While the workflow is robust across diverse datasets, these factors highlight the importance of consistent acquisition settings and careful data preparation when applying the pipeline to new imaging systems or experimental conditions. Beat rate detection not infallible; however, a manual override option is provided to allow user-guided correction when required.

## CONCLUSION

We developed a Python-based analysis framework for label-free cardiac imaging that enables automated extraction of contraction waveforms, speed traces, and wave-propagation dynamics from transmitted light time-series data. The pipeline integrates seamlessly with TIFF-based inputs, supports both single-file and batch processing, and provides calibrated, biologically interpretable outputs. Implemented in a Jupyter notebook with an intuitive interface, it reproduces all analytical steps of the original MATLAB workflow while improving accessibility and scalability. The framework is robust across datasets, requiring only minimal parameter adjustments and no structural code changes.

This work establishes a generalisable pipeline for analysing label-free cardiac timelapse data across diverse experimental systems. By demonstrating consistent performance in hiPSC-CM cultures, killifish recordings, and independent public datasets, the framework shows clear applicability beyond a single laboratory or imaging platform. Its open, notebook-based implementation supports transparent parameter control, straightforward reuse, and integration with community datasets, strengthening reproducibility in functional imaging research. Together, these features position the pipeline as a practical and accessible tool for standardising cardiac motion analysis across labs and repositories.

## ACKNOWLEDGEMENTS

RABB received research funds from the Ellis T. Davies Fellowship Endowment, University of Liverpool. We acknowledge Animal Free Research UK (AFRUK) for the experiments performed without animal products (differentiated hiPSC-CMs on human fibronectin). AD is funded by J Hill Abram Prize PhD studentship at the University of Liverpool awarded to RABB. We acknowledge the Liverpool Centre for Cell Imaging (CCI) for provision of imaging equipment and technical assistance and the Digital Research Infrastructure team for provision of data storage and technical assistance.

## FUNDING

RABB received research funds from the Ellis T. Davies Fellowship Endowment, University of Liverpool. We acknowledge Animal Free Research UK Grant (Developing more realistic human-based cell culture models to study cardiac arrhythmias) for the experiments performed without animal products (hiPSC-CMs). The Cell Discoverer 7 was funded by MRC grant number MR/X013502/1. AC acknowledges funding from the Royal Society, grant number IF\R2\242142.

## CONFLICT OF INTERESTS

RABB is an editorial board member on Scientific Reports. The other authors declare no competing interests.

## DISCLAMERS

None

## AUTHOR CONTRIBUTIONS

RABB conceived and designed research; WP, TW, BB, AD and RABB performed experiments; PS provided hiPSC-CMs; RH and TK collected killifish data; AC first wrote early Matlab scripts; WP, RABB, AC, MH interpreted results of experiments; MH wrote the python script and developed the code; WP, RABB, MH drafted manuscript; WP, MH, AC and RABB edited and revised manuscript; All authors approved final version of manuscript.

## DATA AVAILABILITY

See ’Availability of materials and data’ section for more information.

## MATERIALS AND METHODS

### Cell Culture

The iPSC differentiation process followed protocols described in earlier studies^36,37^. In summary, undifferentiated, healthy human stem cells (ATCC) were plated on Geltrex-coated culture dishes at a density of 1 × 10^5 cells/cm^2^ and cultured for 48 hours until reaching approximately 80% confluency. Differentiation was initiated (day 1) by replacing the medium with StemPro-34 (Life Technologies, 10639011) supplemented with recombinant human Activin A (Life Technologies, PHC9564) and recombinant human BMP4 (R&D Systems, 314-BP-010) for 48 hours. On day 3, the culture medium was switched to RPMI (Life Technologies, 61870036) containing 1× B-27 supplement (Life Technologies, 17504044), the small molecule inhibitor KY02111 (Bio-Techne, 4731/5), and XAV939 (Bio-Techne, 3748/10). From day 5 onward, cells were maintained in RPMI with B-27 supplement only, with medium changes every other day. Human iPSC-cardiomyocytes (hiPSC-CMs) were plated on 96-well dishes at 1 x 10^5^ cells/well. Wells were pretreated with Geltrex™ (A1413302, Thermo-Fisher Scientific) at 0.5μg/cm2, incubated for 1 hour at 37°C and then at room temperature for 1 hour, or human fibronectin (animal free product) used at 2.5 µg/cm^2^ (Catalogue number 11051407001 Roche, from Sigma). In brief, for human fibronectin, the working concentration was 5 μg/cm^2^ for the coating of cell culture vessels. Add 1- or 5-mL sterile water, respectively, to yield a final concentration of 1 mg/mL. Incubate 30 to 60 minutes at 37 °C to dissolve. The culture dish surfaces were coated with human fibronectin under sterile conditions by applying a minimal volume sufficient to cover the entire area. Coated surfaces were then air-dried at room temperature in a tissue culture hood for at least 45 minutes before seeding cells and adding culture medium.

3D heterocellular cardiac spheroids were produced in ultra-low attachment 96-well plates (Corning) as previously described^38^. Human embryonic stem cell-derived cardiomyocytes, human cardiac fibroblasts, and human cardiac endothelial cells were combined at a 4:2:1 ratio, with 10,000 cardiomyocytes seeded per spheroid. Cultures were maintained in a 1:1 mixture of RPMI/B-27 and endothelial growth medium (MV2), and spheroids were allowed to mature for 2-3 weeks before analysis. Fibroblast and endothelial cell cultures were used up to passage 9. The resulting spheroids displayed spontaneous beating behaviour throughout the culture period.

### Imaging

All imaging was performed on a CD7 (Carl Zeiss Microscopy GmbH, Germany) with Zen Blue (3.7) software at the Centre for Cell Imaging (CCI, Liverpool). The 96-well plate with 1.0 x 10^5^ CMs/well was imaged once the cells were steadily beating (start beating 8-10 days after differentiation and imaged 3-5 days once the cells started beating). The environmental conditions were controlled at 37°C and with a supply of 5% CO_2_. Sequential well imaging was set up manually, as automatic imaging of multiple wells requires autofocus, which is incompatible with the CD7’s fast camera streaming.

Time-series imaging of hiPSC-CM monolayers and cardiac spheroids was performed using a Zeiss CD 7 microscope equipped with a white-light LED illumination source, a Plan-Achromat 5×/0.35 NA objective and a Zeiss Axiocam 712 mono camera (Table 2). A 0.5× tube lens was used, giving an effective numerical aperture of 0.125 and a reported depth of focus of 8.98 µm. Images were acquired using transmitted-light brightfield contrast. The optical configuration included filters with wavelength ranges of 412–438, 501–527, 583–601 and 662–756 nm, and beam splitters at 394, 490, 573 and 691 nm.

For hiPSC-CM monolayer time series, the white-light LED illumination intensity was set to 1 %. Images were acquired with a 629 µs exposure time at 8-bit depth, without binning (1 × 1) for a total of 19.77 sec (612 frames). For cardiac spheroid time series, the illumination intensity was set to 12 %, with an exposure time of 200 µs and an image bit depth of 14 bits, also without binning (1 × 1) for a total of 19.69 sec (2530 frames). Both acquisition configurations used a spatial sampling of 1.38 µm per pixel. The resulting image sequences were subsequently analysed using the BeatRatePy pipeline.

### Datasets

**Table 3:** Overview of the image data used in this study. Datasets and their respective metadata relevant for the processing. Metadata from data type “own” was extracted from the proprietary microscopy files. For public datasets, the metadata was extracted either from the publication/supplementary materials and/or the downloaded video files via FFMPEG. Any video files were converted into a tiff sequence using FFMPEG, imported into Fiji, and converted into an 8-bit grey scale time series. The spatial calibration of the images was generally not obtainable.

| ID | Name | Datab<br>ase | Accession<br>number/file<br>name | Sample type | Reference |
| --- | --- | --- | --- | --- | --- |
| 1 | Cardiac<br>cell<br>culture | Biolma<br>ge<br>Archiv<br>e | S-BIAD4119 | Cardiac cell culture | https://doi.org/10.6019/S-BIAD4119 |
| 2 | Cardiac<br>spheroid | Biolma<br>ge<br>Archiv<br>e | S-BIAD4119 | Cardiac spheroid | https://doi.org/10.6019/S-BIAD4119 |
| 3 | Killifish | Univer<br>sity of<br>Exeter | https://doi.org/10.24378/exe.5506<br>Files used:<br>Submission<br>data\Cropp<br>ed<br>greyscale<br>frames<br>(TIFFs)\16<br>file: 16-1-R | Killifish embryo heart | https://doi.org/10.1364/BOE.578208 |
| 4 | Muscle<br>Motion | N/A | Publication<br>supplement<br>al Material<br>312067_onl<br>ine_movie_<br>iii.tif | Human stem cell<br>derived<br>cardiomyocyte<br>monolayer culture | Muscle Motion:<br>https://doi.org/10.1161/CIRCRESAHA.117.312067 |
| 5 | Muscle<br>Motion | N/A | Publication<br>supplement<br>al Material<br>312067_onl<br>ine_movie_<br>iv.tif | Human stem cell<br>derived cardiac<br>organoid. | Muscle Motion:<br>https://doi.org/10.1161/CIRCRESAHA.117.312067 |
| 6 | Muscle<br>Motion | N/A | Publication<br>supplement<br>al Material<br>312067_onl<br>ine_movie_<br>xi.tif | Echocardiogram of a<br>healthy individual | Muscle Motion:<br>https://doi.org/10.1161/CIRCRESAHA.117.312067 |
| 7 | Cardiac<br>Wave | N/A | hiPSC-CM_CTR.avi | Human-induced<br>pluripotent stem<br>cell-derived<br>cardiomyocytes in<br>culture | Dense optical flow software to<br>quantify cellular contractility:<br>https://doi.org/10.1016/j.crmeth.2021.100044 |
| 8 | Cardiac<br>Wave | N/A | hiPSC-CM_ISO.avi | Drug treated human-induced pluripotent<br>stem cell-derived | Dense optical flow software to<br>quantify cellular contractility: |
|  |  |  |  | cardiomyocytes in culture | https://doi.org/10.1016/j.crmeth.2021.100044 |
| 9 | Cardiac Wave | N/A | ADULT-CM_CTR.avi | Acutely isolated adult mice or rat ventricular myocytes | Dense optical flow software to quantify cellular contractility: https://doi.org/10.1016/j.crmeth.2021.100044 |
| 10 | Cardiac Wave | N/A | ADULT-CM_ISO.avi | Drug treated acutely isolated adult mice or rat ventricular myocytes | Dense optical flow software to quantify cellular contractility: https://doi.org/10.1016/j.crmeth.2021.100044 |

### Availability of materials and data

The initial script by Dr Alex Corbett (https://gitlab.com/psfcheck/pixel-variance/-/blob/main/PixelVariance_Matlab?ref_type=heads) was written in MATLAB and translated by Dr Marie Held into Python. The code, maintaining all processing and analysis steps, was implemented in python utilising the freely available tifffile, bioio, numpy, scipy and matplotlib packages. The graphical user interface uses jupyter lab, ipywidgets and napari. Figures are made using matplotlib and jupyter lab. An up-to-date version of the code is available on Github together with information on how to install and run the notebook locally: https://github.com/Marien-kaefer/BeatRatePy

Our algorithm offers several advantages over existing tools for quantifying cardiac contraction from label-free imaging (Table 4). BeatRatePy is freely available through a Jupyter notebook, a web-based interface that combines code, narrative text, equations and rich output types in a simple interface accessible to users with limited coding experience. The pipeline uses a pixel variance-based algorithm, supports batch processing with automated outputs, and works natively with proprietary microscopy file formats without requiring file conversion. It automatically calculates beat rate alongside further outputs, and can analyse both cell monolayers and individual cells or spheroids. By contrast, other tools such as Myocyter and HeartCV require AVI video files as input, which may have undergone lossy compression – for example, when recordings are made using an iPhone, as in Myocyter – although the codecs used are not always reported; BeatRatePy instead processes raw microscopy data directly, avoiding this potential source of error. The pipeline is also extensible, allowing further functionality to be added in future.

**Table 4:** Feature comparison between BeatRatePy and existing cardiac motion analysis tools. Comparison of BeatRatePy with commonly used tools for label-free cardiac motion analysis, including MuscleMotion, Myocyter, and ContractionWave. Features assessed include software accessibility, compatibility with proprietary microscopy formats, batch-processing capability, spatial calibration requirements, and automated beat-rate determination. BeatRatePy combines native support for proprietary microscopy data formats with automated beat-rate quantification, optional spatial calibration, and batch processing within a user-friendly Jupyter notebook environment, facilitating reproducible analysis across a range of cardiac imaging datasets.

| Feature | "BeatRatePy" | MuscleMotion | Myocyter | Contraction Wave |
| --- | --- | --- | --- | --- |
| Free to use software | Yes | Yes | Yes | Yes |
| Proprietary microscopy format compatibility | Yes | Yes | No | No |
| Batch processing | Yes | No | No | Yes |
| Spatial Calibration | Optional | No | No | Mandatory |
| Automated beat rate output | Yes | No | Yes (reported as frequency) | No |

### Data Input

The “BeatRatePy” algorithm operates on TIFF files composed of a time series of images. The jupyter notebook requires the following adjustable inputs (ref to jupyter notebook figure):

1. Folder in which the file(s) is/are located.
2. Checkbox to confirm batch processing of all imaging files located in the folder specified in 1.
3. Filename – only required if batch processing checkbox (2.) is unchecked.
4. Video rate in frames per second.
5. Checkbox to confirm the automated calculation of the beat rate from the input time series.
6. Manually determined Beat rate in Hz – only required if automated beat rate calculation (5.) is unchecked
7. Offset in frames – only required if calculate offset from video & beat rate checkbox (8.) is checked.
8. Checkbox to confirm calculation of the offset (frames) from the video rate, beat rate and fractional offset.
9. Fractional offset for automated offset calculation.
10. Checkbox to confirm calibration of the outputs with the pixel calibration.
11. Pixel calibration in micrometre – only required if checkbox in 10 is ticked.

Furthermore, at least one time series file is required. For batch processing, folders containing the multiple image files recorded at the same frame rate and pixel calibration (if used for calculation) are required.

### Data Pre-processing

For CD7-generated data, following image collection in ZEN. The video rate and pixel calibration were the same for each time series and they could therefore be processed in batch but would need to be considered in the sorting of data from different microscopes or imaging settings.

For data provided as video files, the videos were first converted from video format (AVI, MOV, MP4) to TIFF sequences through FFMPEG [ffmpeg tool (Version 2025-03-24-git-cbbc927a67-full_build- www.gyan.dev). Available from http://ffmpeg.org/]. FFMPEG is a free and open-source software which during conversion also displays the frame rate for the video file, which was recorded.

The tiff sequence was subsequently imported into Fiji^39^, converted from RGB to 8bit grey scale, re-ordered as a time series, the frame rate added and saved as a time-lapse tiff stack. Where required, relevant cropped regions were generated. Where multiple spheroids were present in the same field of view, individual spheroids were cropped prior to analysis.

### Data viewer

Whilst the TIFF file format is compatible with many data viewers, we chose the Napari viewer for convenience. For single datasets being processed the raw data were imported and displayed as an image layer in napari viewer. Following the calculation of the pixel variance, this data, including a padding of several black frames equating to the number of offset frames, will also be displayed as a second image layer in napari (Supplement Figure 1). For batch processing, data is not visualised and cleared from memory after each file has been processed to optimise system performance.

### Time Series Analysis

BeatRatePy was developed to quantify beat rate, motion (speed), and deformation (contraction) profiles from cardiac imaging datasets. Image stacks were imported and converted to 16-bit arrays. Beat rate was estimated from the mean absolute pixel intensity difference between consecutive frames to generate a motion trace. This trace was detrended by subtracting a moving-average baseline using a window equal to 10% of the total trace length and subsequently smoothed using a Savitzky-Golay filter (window size = 11 frames, polynomial order = 3). Beat rate was determined by autocorrelation analysis of the processed motion trace, restricting the analysis to lags greater than 0.3 s to exclude the zero-lag peak and physiologically implausible frequencies exceeding 200 beats per minute. The lag corresponding to the dominant autocorrelation peak was converted to a beat period using the acquisition frame rate, from which beat rate was calculated and reported in both hertz (Hz) and beats per minute (BPM).

Motion (speed) and deformation (contraction) profiles were quantified using the pixel-variance approach described previously^31^. A rolling pixel variance was calculated over a frame offset determined from the acquisition frame rate and calculated beat frequency.

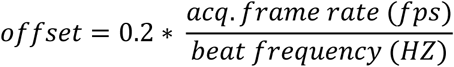

The resulting variance stack was converted from a 32-bit to a 16-bit range, zero-padded with several frames equivalent to the frame offset, and saved as a TIFF stack. Mean pixel intensities from each variance image were used to generate a speed trace, which could be optionally scaled to µm/s when pixel calibration data were available. The frame exhibiting minimal speed was identified as the key frame, and the absolute difference in pixel variance relative to this frame was calculated for each image to generate a contraction trace, optionally scaled to µm. Speed and contraction traces were smoothed using a Savitzky-Golay filter (window = 11 frames, order = 3), normalised, plotted, and exported together with the corresponding scaled measurements.

### Output

A subfolder called “analysis” will be created automatically in the specified input folder containing the time-lapse data. The ten chosen input parameters and options are saved as a text file for accountability and reproducibility purposes. The generated output files comprise the pixel variance ome.tiff stack, a speed text file containing three columns (time, speed in px/s or μm/s, min-max normalised speed) and contraction text file containing three columns (time, contraction in px or μm, min-max normalised contraction), as well as plotted speed or contraction over time, both absolute and normalised values. The applied file naming convention preserves the original file name and suffixes each output file appropriately.

### Figure Creation

Multipaneled microscopy figures for this publication were created with OMERO.figure, a web-based Open Microscopy Environment (OME) application designed for rapid generation of scientific figures for publications and presentations directly from image data stored within an OMERO database^40^ together with Inkscape for figure assembly and annotation (https://inkscape.org/).

### Beat Rate Extraction from Other Packages

MUSCLEMOTION was run in Fiji (1.54p, stable version), the manually measured beat rate was used to calculate the speed Window from the beat rate, video frame rate and an offset fraction of 0.2. The video frame rate was specified in the Muscle Motion opening dialogue accordingly (Supplemental Table 1) and otherwise the default settings were used. The beat rate was calculated from the Log_file.txt by using the list of Peaks detected at points (frames) and the frame rate.

Myocyter was run in Fiji version (2.0.0-rc-69/1.52n) as recommended in the original publication and the beat rate is an output parameter that is written into the results text file.

In ContractionWave, the beat frequency can be output by selecting the trace or part thereof followed by clicking on the Fast Fourier Transform option. For both packages, the beat rates were calculated/extracted for each input file individually.

## SUPPLEMENTAL MATERIAL

**Supplement Table 1:**
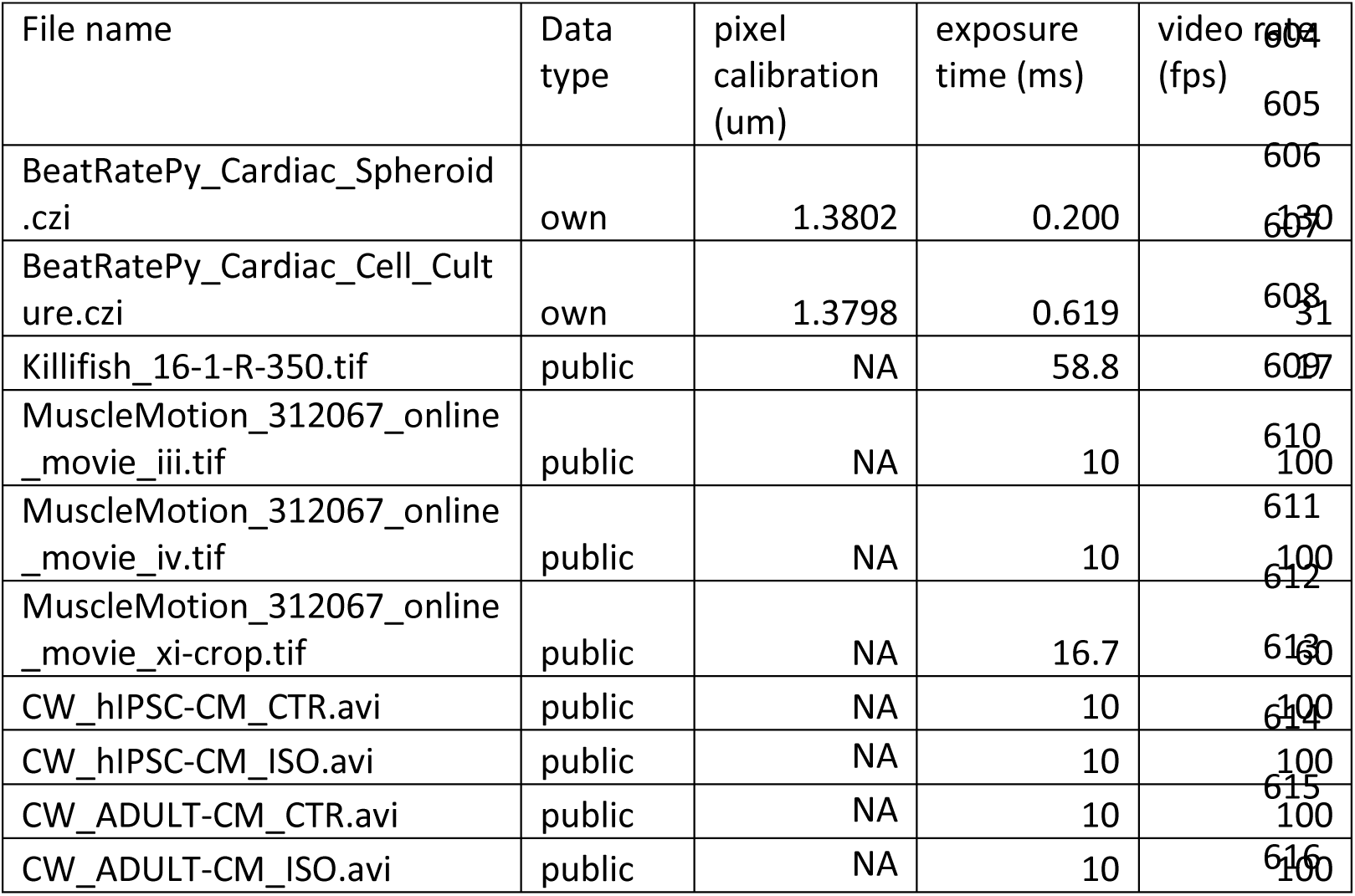
Summary of the imaging datasets analysed in this study, including data source (own or public), pixel calibration, exposure time, and video acquisition rate used for BeatRatePy processing and analysis.

**Supplement Figure 1:**
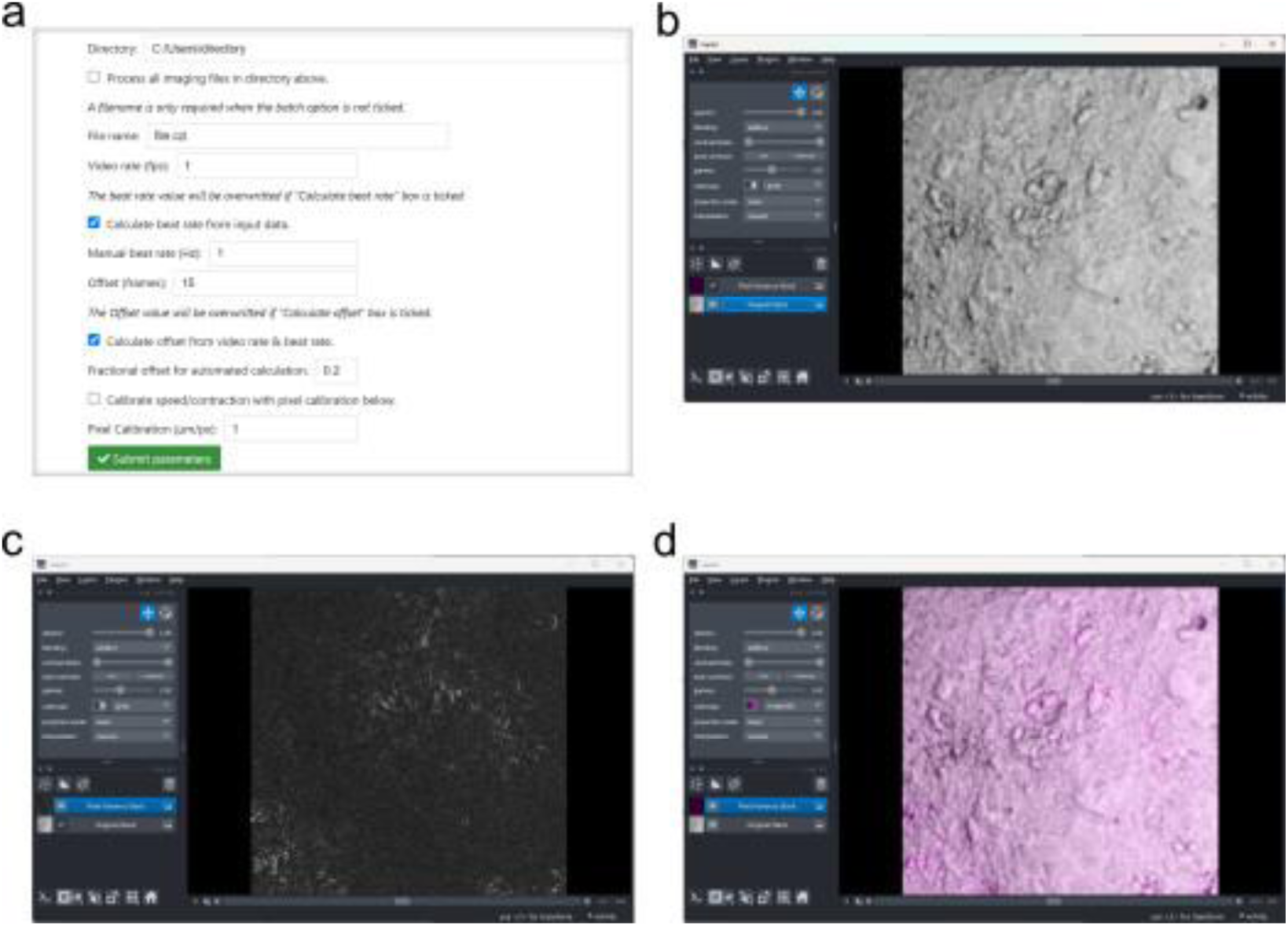
Graphical workflow of the BeatRatePy pipeline. The Jupyter notebook interface (a) provides user-adjustable parameters for image analysis. Original image data are visualised in Napari (b), followed by generation of a pixel variance image stack highlighting regions of motion (c). A merged view of the original grayscale images and pixel variance signal (magenta) is displayed (d), facilitating visual validation of motion detection prior to extraction of speed, contraction, and beat-rate measurements.

### Supplemental Videos

1. BeatRatePy_SM_Cell_culture_full_field_of_view.mp4 - Live imaging of a contracting monolayer of hiPSC-Cardiomyocytes

Video showing a multi panel time series consisting of a transmitted light (left), pixel variance (centre) and merged image panel (right) of a full field of view. The sequence of images was rendered at 31fps showing recorded data in “real time”. The transmitted light sequence was pre-processed with a rolling ball background subtraction filter to remove the microscope objective’s vignetting effect for visual purposes.

2. BeatRatePY_SM_Cell_Culture_crop.mp4 – Detail of live imaging of a contracting monolayer of hiPSC-Cardiomyocytes

Video showing a multi panel time series consisting of transmitted light (left), pixel variance (centre) and merged image panels (right) of a zoomed in section of the full field of view recorded and shown in Supplemental video 1. The sequence of images was rendered at 31fps showing recorded data in “real time”.

3. BeatRatePy_SM_Spheroid.mp4 - Live imaging of a contracting heterocellular cardiac spheroid

Video showing a multi panel time series consisting of transmitted light (left), pixel variance (centre) and merged image panels (right). The sequence of images was rendered at 130fps showing recorded data in “real time”.

## Notes

https://doi.org/10.6019/S-BIAD4119

https://doi.org/10.1364/BOE.578208

### Summary of Updates

The title had a typo " Hight-Throughput..."

